# Programmatic access to ICTV virus taxonomy through a public ontology API

**DOI:** 10.64898/2026.06.16.732600

**Authors:** Philippe Lieutaud, James McLaughlin, R. Curtis Hendrickson, Romain David, Helen Parkinson, Elliot J. Lefkowitz, Donald M. Dempsey, Bruno Coutard

## Abstract

The International Committee on Taxonomy of Viruses (ICTV) is responsible for developing and maintaining a universal virus taxonomy. As the reference framework for organising the viral world, it is essential for virology and related fields. Despite its widespread use in research and public health, programmatic access to ICTV taxonomy has remained limited, posing challenges for integration, versioning, and interoperability across databases and bioinformatics resources requiring up-to-date virus taxonomy. To address this, we developed a public and sustainable solution leveraging ontology-based APIs. Successive ICTV Master Species List (MSL) releases were transformed into a structured ontology and deployed as a unified representation through the Ontology Lookup Service (OLS). The framework also provides ICTV–NCBI mappings and helper libraries for integration into downstream systems. This enables, for the first time, public programmatic retrieval of current and historical virological taxon names, taxonomic relationships, metadata, and persistent identifiers through stable endpoints. More broadly, this work illustrates a general strategy for transforming structured biological datasets into semantically enriched graph resources exposed through scalable public APIs. These developments enhance interoperability, reduce manual curation, and support FAIR-aligned taxonomic data management in virology and pandemic preparedness.

**Key points:**

- ICTV provides the official taxonomy for classifying viruses and naming virus taxa, but lacks standardised programmatic access.
- Transforming ICTV data into an ontology enables semantic, machine-actionable access across releases via ontology-based APIs.
- ICTV–NCBI mappings support interoperability across bioinformatics resources.
- The framework enables programmatic resolution of current and historical viral taxa.
- This approach provides a reusable model for exposing biological datasets through public APIs.

## 1 Introduction and landscape

The [International Committee on Taxonomy of Viruses (ICTV)](https://ictv.global/), established under the Virology Division of the [International Union of Microbiological Societies (IUMS)](https://www.iums.org/), is the authoritative body responsible for the classification of viruses and naming of virus taxa, ensuring a standardised and commonly accepted nomenclature. Virus taxonomy provides a structured framework for classifying viruses based on evolutionary relationships, shared biological properties, and genomic features, such as genetic material, structural features, replication strategies and host range. While everyday virus names function as vernacular labels that can vary across studies and contexts, the ICTV taxon names provide formal, standardised designations that ensure unambiguous communication across the scientific community [1].

The regularly updated ICTV Master Species Lists (MSLs) reflect ongoing advances in virology, including classification of newly discovered viruses and refined taxonomic insights, with one major new release each year, sometimes complemented by minor releases [2]. As virus discovery accelerates, particularly through metagenomics, maintaining consistent and up-to-date taxonomic references in databases and bioinformatics tools relying on ICTV taxa becomes increasingly important.

While the ICTV releases provide authoritative reference data, their distribution format is primarily designed for human consultation and does not readily support programmatic access [3]. As a result, integrating ICTV taxonomy updates into third-party computational systems remains challenging. Broader taxonomic platforms such as [NCBI Taxonomy (National Center for Biotechnology Information)](https://www.ncbi.nlm.nih.gov/), [GBIF (Global Biodiversity Information Facility)](https://www.gbif.org/), [COL (Catalogue of Life)](https://www.catalogueoflife.org/) and Wikidata embed ICTV releases as the authoritative source for virus taxonomy. However, these platforms often incorporate ICTV updates with substantial delays and may lack systematic support for resolving historical taxonomic changes.

Downstream resources can lag by one or more ICTV taxonomic updates, resulting in discrepancies in the classification they expose. Such inconsistencies create significant challenges for users who require the most up-to-date virus classifications, potentially affecting research accuracy and data consistency. For example, outdated species names may persist for some time in major sequence repositories such as [GenBank](https://www.ncbi.nlm.nih.gov/genbank/) and the [European Nucleotide Archive (ENA)](https://www.ebi.ac.uk/ena/browser/home), leading to further propagation of downstream inconsistencies. Since these repositories rely on the NCBI Taxonomy as their primary taxonomic reference, such delays can be observed by comparing historical taxonomic archives of NCBI and ICTV. A concrete example is ICTV renaming “*Severe acute respiratory syndrome-related coronavirus”* to”*Betacoronavirus pandemicum”* in MSL39. This change did not appear in NCBI Taxonomy for more than a year, and, when it did, was represented by the creation of a new NCBI taxon, rather than a direct renaming of the existing taxa. Obsolete taxon names that continue to be treated as current references complicate reproducibility and hinder interoperability across resources. These problems are inevitable, given that NCBI is not simply embedding ICTV taxonomy, but integrating it within a broader system with additional constraints. As such, ICTV remains the authoritative source for virus taxonomy, while NCBI serves as a widely used integration layer. Such cases illustrate why programmatic resolution mechanisms are needed [4], along with traceable ICTV identifiers.

The [European Viral Outbreak Response Alliance (EVORA)](https://evora-project.eu/) project is a European initiative (2024-2026; grant No. 101131959) that aims to enhance pandemic preparedness and response by strengthening research coordination, regulatory alignment, and data sharing among EU Research Infrastructures. Improving universal adoption of accurate and up-to-date virus taxonomy represents an important component of this mission. Members of EVORA and ICTV collaborated to provide a structured, machine-actionable version of the official ICTV taxonomy using ontology-based APIs, with support for historical name resolution, revision tracking and cross-system identifiers.

Many biological data resources are distributed as relational databases, spreadsheets or tabular datasets that are well suited for storage, but less adapted to programmatic integration across heterogeneous systems. In addition to addressing the immediate need for improved access to ICTV taxonomy, this work illustrates a broader methodological approach for transforming structured database content into semantically enriched knowledge representations through the use of ontologies. Ontologies play a crucial role in the management, integration, and analysis of biological data, providing a structured framework to annotate, share, and interpret complex datasets and to make meaning human and machine readable [5]. Converting ICTV releases into an ontology enables graph-based querying and integration. As graph-based representations naturally capture hierarchical relationships, synonyms, and historical changes, representing taxonomy as an ontology therefore enables more expressive queries, automated reasoning, and easier integration with other semantic resources.

## 2 Methods and strategy

In this paper, “OLS API” refers to the generic programmatic interface provided by the OLS, including its Representational State Transfer (REST) endpoints and controllers. “ICTV ontology API” refers to access to the ICTV ontology through the OLS API, either directly via OLS endpoints or indirectly through ICTV-specific helper libraries built on top of them.

### 2.1 Design requirements and use cases

To support practical integration of ICTV taxonomy into data systems and analytical workflows, several key use cases were defined to address common end-user challenges for automated processing. First, it must provide historical resolution by mapping obsolete names, synonyms, and ICTV identifiers from previous ICTV releases to their corresponding taxa in the current ICTV classification. Second, taxonomic changes must be tracked across releases, including taxa that have been abolished, split, merged, renamed, or reassigned (promoted) within the hierarchy. Third, cross-resource mapping must be supported, and, specifically, a mapping, with provenance, between NCBI Taxonomy identifiers and ICTV identifiers (historical and current) must be provided. Finally, there must be a stable, public, well-documented API for performing these functions.

### 2.2 Persistent identifiers

Persistent identifiers are key for data integration [6]. The ICTV defines persistent identifiers for all versions of all taxa, as well as links connecting the taxa into hierarchies and documenting changes between releases. These unique identifiers facilitate resolution of outdated taxonomic terms to current ones by allowing taxa to be efficiently tracked across renames, rank changes, merges, splits, and abolitions between releases. To support programmatic resolution and cross-release linkage, and to further support FAIR principles, ICTV publicly documented these existing identifiers and defined Compact Uniform Resource Identifiers (CURIEs) for them. ICTV provides three principal identifiers. In the shorthand used below, the letter prefix indicates the identifier type and # represents the numeric identifier value:

- Taxnode identifier (Taxnode ID; TN#): uniquely identifies a taxon in a specific release.
- ICTV identifier (ICTV ID; ICTV#): identifies the same taxon across releases, including through renames, changes in location within the taxonomic tree (moves), and changes in rank (promotion/demotion). Splits and merges create new ICTV IDs, which are linked back to their antecedent taxa.
- Isolate identifier (Isolate ID; VMR#): defined by the Virus Metadata Resource (VMR), links GenBank sequence accessions for one or more segments to the vernacular name and the corresponding viral species.

These identifiers provided the basis for modelling ICTV releases as semantically linked ontology resources. An extremely abridged example of the history of a taxon related to the genus *Baculovirus* in Figure 1 illustrates some of the complexity, and demonstrates how ICTV IDs link historic versions of a taxon, from the first ICTV release to the current MSL41 release [7].

**Figure 1:**
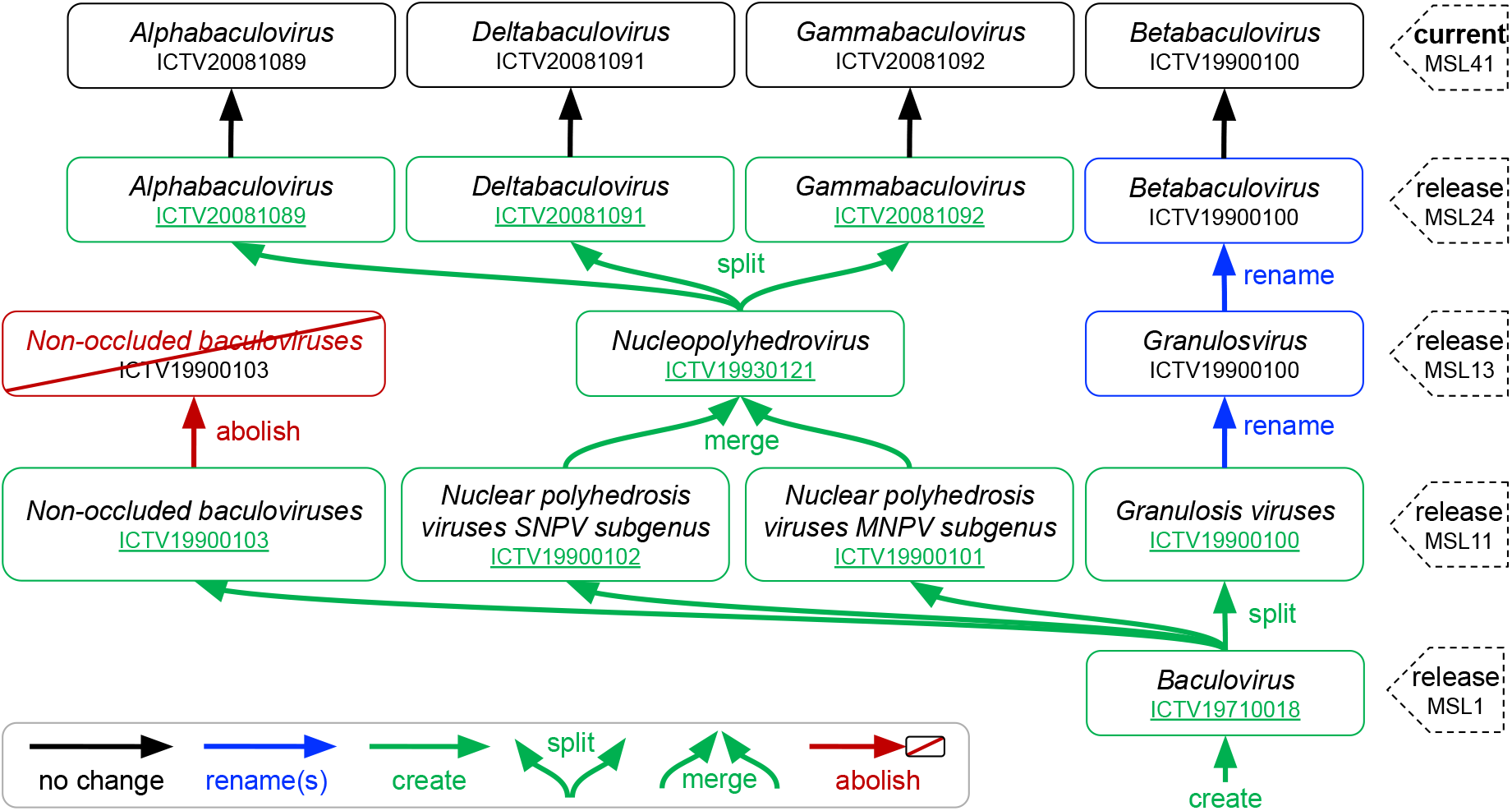
Illustrative example of taxonomic changes across ICTV releases using a genus-level taxon first recorded in MSL1. The diagram shows historical and current genus names originating with the genus *Baculovirus* (ICTV19710018), first recorded in the first ICTV release, and ending with the four currently recognised genera. This example illustrates how the grouping of species into genera, although species are not shown here, can be redefined over time as viruses are better understood and characterised. In this historical trajectory, the original genus was first split into four genera; two of these were then merged; and the resulting lineage was later split into three genera. This illustrates some of the complexity that can occur in the history of a single taxon, and more broadly how taxonomic changes across different ranks and releases can make the overall taxonomy difficult to trace. It also shows how ICTV IDs are assigned to link historical versions of the same taxon. When a taxon is created, split or merged, indicated by green single, diverging or converging arrows, respectively, a new ICTV ID is assigned and shown underlined in green. When taxa are renamed, promoted, demoted or moved, the ICTV ID remains the same, linking the different versions. **Alt text:** Diagram showing historical and current ICTV genus names and identifiers originating with *Baculovirus* (ICTV19710018) and ending with four currently recognised genera. Green arrows indicate taxon creation, splits and merges across releases.

### 2.3 Initial modelling of ICTV releases as ontologies

To meet the design requirements and use cases described above, including historical resolution, cross-release tracking and programmatic querying, each of the 41 (as of 2026) official ICTV taxonomy updates since 1971 was modelled as an independent OWL2 ontology, representing hierarchies, ranks, synonyms and revision metadata. Taxa were modelled as classes and virus isolates as individuals. This approach captures taxonomic relationships and their evolution over time, while preserving the structural and semantic fidelity of the original ICTV data in a machine-readable format.

The ontologies were constructed using established semantic standards and interoperable vocabularies commonly used in biomedical ontology ecosystems (e.g. [8]), built upon semantic web standards including OWL, RDFS, PROV-O [9] and SKOS semantic relations [10], together with complementary vocabularies such as Dublin Core Terms (DCTERMS), Friend Of A Friend (FOAF), the Taxonomic Rank Vocabulary (TAXRANK) and oboInOwl ontology annotations. This ensures compatibility with the broader life-science ontology ecosystem [11] and consistency with OBO Foundry principles [12]. Obsolete taxa were linked to their replacements using IAO:0100001 (term replaced by), enabling rapid resolution of obsolete terms into their current versions.

The Ontology Lookup Service (OLS) enables direct exposure of ontology data through stable REST API controllers, a web-based user interface [13], and a Model Context Protocol (MCP) server for access by AI agents. OLS provides a ready-to-use infrastructure to host ontologies and deliver programmatic access through standardised endpoints [14]. This approach eliminates the need to develop and maintain custom API infrastructures while ensuring compatibility with the broader ecosystem of life science ontologies. Thus, transforming existing data into a semantically structured graph representation leverages the OLS infrastructure to efficiently provide standard, public APIs and interoperability across data infrastructures.

The set of release-specific ontologies was published via OLS. The OLS API enables dynamic querying, retrieval of data and linking across ontologies, allowing applications to access up-to-date information directly from ICTV data via the ontology stored in OLS.

However, querying multiple ontologies proved complex for the primary use case of reconciling historical references with the current taxonomy. The system was therefore redesigned to merge all releases into a single, unified ontology (over 195,000 terms in 2026). In this model, the most recent ICTV release represents the authoritative classification, while entities from previous releases are preserved as obsolete terms linked through explicit replacement relations. This design reduces query complexity, while preserving full history, provenance and backward compatibility across ICTV releases. This enables any taxonomic reference appearing in historical datasets or publications to be programmatically reconciled with the current ICTV classification.

### 2.4 Availability and sustainability

Continuous Integration and Deployment (CI/CD) workflows implemented through public GitHub repositories automatically trigger the ontology generation workflow for each release published in the ICTV’s official GitHub repository. This automation removes manual intervention and contributes to long-term sustainability of the infrastructure, in line with recommendations for sustainable FAIR implementation in Life Sciences [15]. The unified ontology is immediately available as a release from the EVORA GitHub repository, and is automatically deployed to the OLS, typically within 24 hours of an ICTV release, which provides public web browsing and API access.

### 2.5 ICTV–NCBI mapping

To improve interoperability at the broader scale [16], ICTV–NCBI mappings are generated using the OLS API, and stored in SSSOM (Simple Standard for Sharing Ontology Mapping) format, aligning ICTV identifiers with NCBI Taxon identifiers [17]. The code to generate this mapping is maintained in a dedicated GitHub repository, where the resulting SSSOM files are then published as releases (see Supplementary Material S4). While this analysis is also automated using a GitHub CI/CD workflow, it runs on a schedule, decoupled from the release cycles of the two ontologies, allowing independent updates while remaining consistent with the continuously updated ontology infrastructure.

These mappings not only facilitate direct and bidirectional linkage between ICTV and NCBI taxonomies, but also enable alignment across downstream resources that rely on NCBI as a taxonomic backbone, including sequence repositories and bioinformatics platforms. This feature enables broader platforms that rely on the NCBI taxonomy for viral references (e.g., ENA) to stay current with the ICTV taxonomy while continuing to leverage the broader coverage of NCBI taxonomy.

The NCBI integrates the ICTV Taxonomy into its own taxonomy system, which is used to designate species related to the entries in its sequence databases (e.g. GenBank). Historically, ICTV updates have been incorporated with roughly a one-year delay, resulting in NCBI being one or two releases behind the latest ICTV version (as described above). Observations following MSL41 release suggest a recent effort by NCBI to shorten this delay, with progressive integration starting in May 2026, about two months after the MSL41 release. This included the integration of renamed taxa, while newly created taxa remained absent and abolished taxa had not yet been removed (see Supplementary Material S3). As part of this work, ICTV has publicly documented several persistent identifiers (described above) that link taxa in successive ICTV Taxonomy versions, which should facilitate rapid integration of new releases. These identifiers are exposed in the ICTV ontology metadata and support the generation of SSSOM ICTV-NCBI mappings. In the released mapping file, lexical ICTV-to-NCBI Taxon correspondences are expressed with skos:exactMatch, enabling alignment between ICTV taxonomy and NCBI Taxonomy identifiers used by downstream sequence resources such as GenBank.

## 3 Results

### 3.0.1 Unified ontology for ICTV taxonomy and OLS deployment

We have produced a unified ontology for the ICTV taxonomy, aggregating all releases into a single resource. The ontology is available for download or via OLS and enables programmatic access to current and historical classifications. Ontology updates are automatically triggered by new ICTV releases and reflected in OLS, typically within 24 hours.

Asynchronously, ICTV–NCBI mappings are generated in SSSOM format, enabling bidirectional resolution between the taxonomies. Mappings are maintained in a dedicated repository and updated independently. The mapping supports bidirectional many-to-many resolution between ICTV and NCBI taxonomies and facilitates integration with downstream resources. The overall architecture of the generation and publication workflow is shown in **Figure 2**.

**Figure 2:**
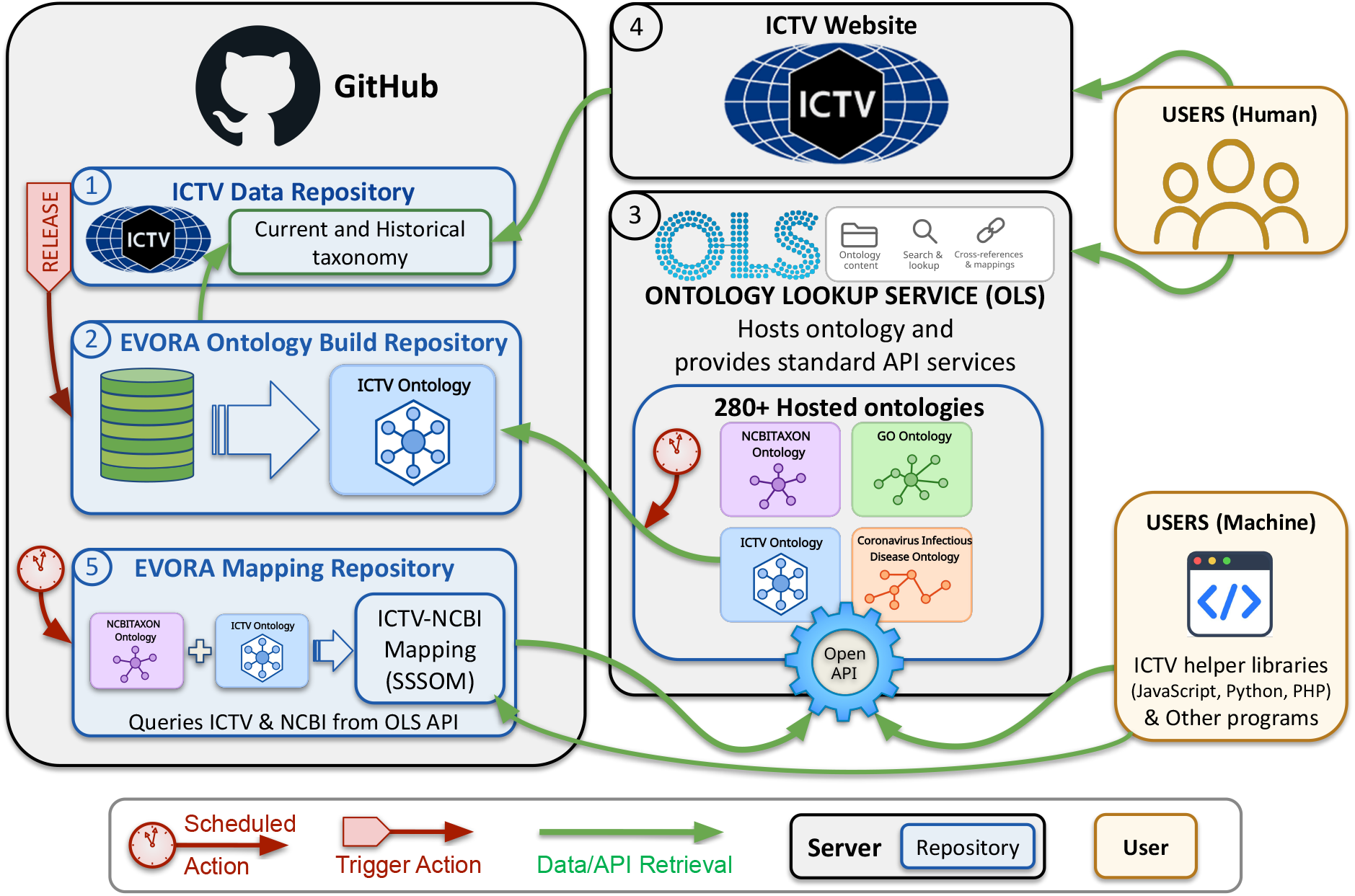
Architecture and workflow of the ICTV ontology and API framework. Official ICTV taxonomy data are released through the public ICTV Data repository (1), hosted on GitHub. A new release triggers the EVORA CI/CD workflow hosted in the EVORA Ontology Build repository (2), which retrieves the ICTV release files, transforms them into OWL ontology artefacts and merges them into a unified ICTV ontology. In this unified ontology, the latest ICTV release represents the current authoritative taxonomy, while entities from previous releases are retained as obsolete terms linked through explicit semantic relations. Through scheduled actions, the unified ontology is loaded into the Ontology Lookup Service (OLS) (3), which provides both a browsable human interface and standard API services. The same ICTV source data also feed the ICTV website (4), which provides the official human-facing access point to ICTV taxonomy. In parallel, a separate EVORA Mapping repository (5) runs scheduled workflows that query OLS to use the latest ICTV and NCBITaxon ontologies to generate ICTV-NCBI mappings in SSSOM format; these mappings are stored in the corresponding public repository. Downstream access is supported through direct API queries and language-specific OLS ICTV API helper libraries, enabling external applications to perform historical taxon resolution, cross-resource mapping and reduced manual curation. Green arrows indicate data/API retrieval actions; peach arrows indicate workflow trigger actions; peach clocks indicate scheduled actions. Black boxes represent servers, blue boxes repositories and brown boxes users. **Alt text:** Workflow diagram showing the end-to-end ICTV ontology and API architecture. Public ICTV taxonomy release data are retrieved by EVORA GitHub workflows, transformed into OWL artefacts, merged into a unified ontology, loaded into OLS for browsing and API access, and used by a separate mapping workflow to generate ICTV-NCBI SSSOM mappings. Symbols distinguish ICTV, EVORA, OLS and user-facing components, including repositories, servers, scheduled actions, workflow triggers and data/API retrieval flows.

To simplify integration into downstream applications, ICTV-specific helper libraries were developed on top of the standard OLS API. These libraries are provided in JavaScript, Python and PHP and encapsulate resolution logic (synonyms, obsolescence, lineage, mappings) while returning normalised data about taxon objects. These objects contain identifiers, labels, synonyms, ranks, lineage, obsolescence metadata and cross-references, enabling application developers to integrate ICTV-aware logic without direct manipulation of raw API calls.

The OLS API documentation, ICTV-specific guidance and [helper libraries are publicly available](https://evora-project.github.io/ictv-ontology/helpers/) (see Supplementary Material S1). Together, these components enable automated resolution, cross-taxonomy linkage and reproducible versioning.

### 3.1 Reference application and early adopters

A public demonstration interface, the ICTV Taxon Resolver web application, was developed as a reference implementation using the JavaScript helper library. The source code is available from the EVORA ICTV Resolver GitHub repository. The resolver allows users to submit current and historical ICTV taxon names and virus names, ICTV identifiers, Internationalized Resource Identifiers (IRIs) or NCBI Taxon IDs and receive the corresponding current, accepted ICTV taxon(s), together with their full lineages and revision history. This tool illustrates practical applicability while serving as a functional community resource. The interface, together with an example resolution of the obsolete species name *Zika virus* to its post-2022 name and lineage, *Orthoflavivirus zikaense*, are shown in **Figure 3**.

**Figure 3:**
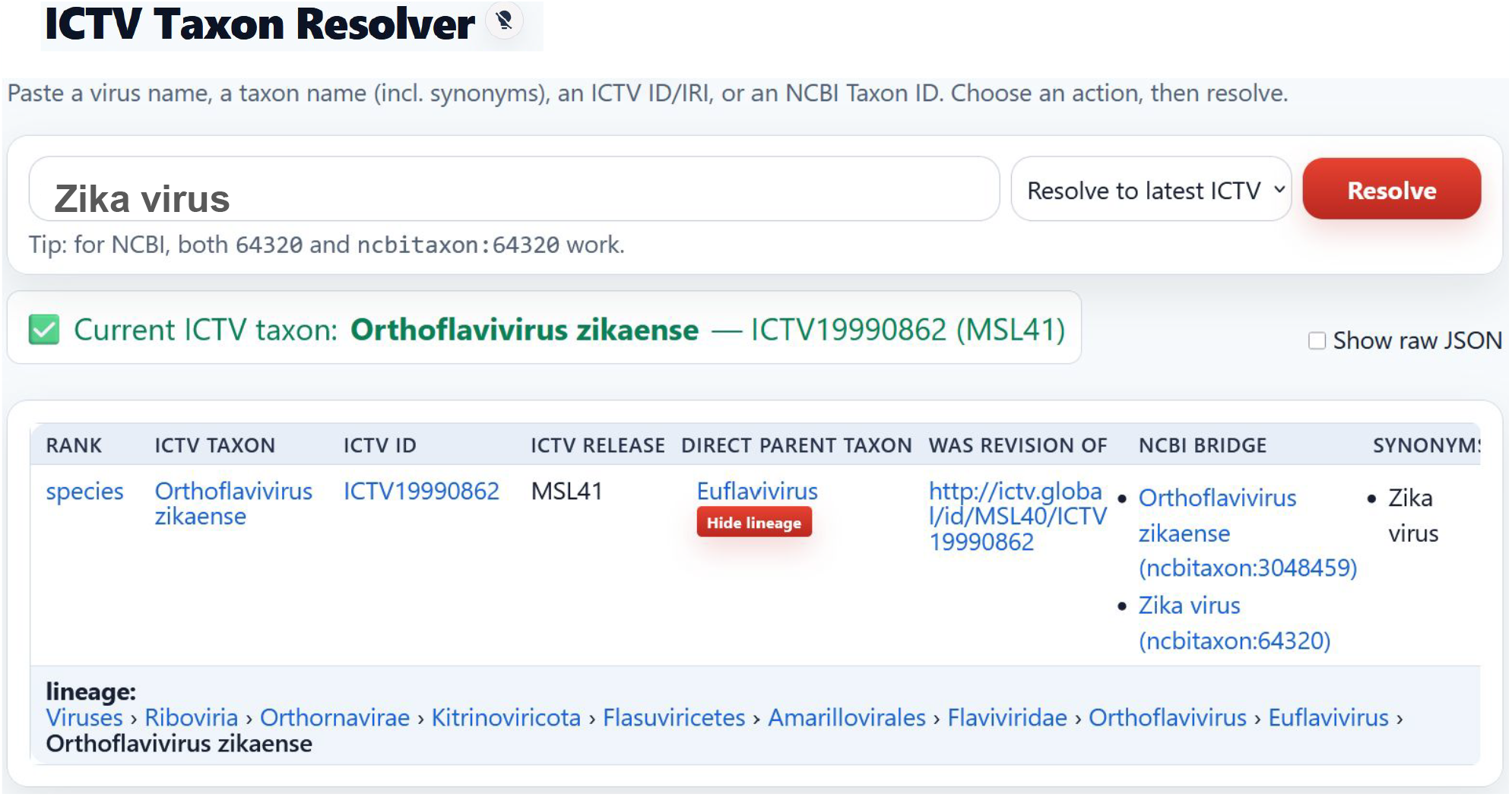
ICTV Taxon Resolver and programmatic taxonomic resolution. The ICTV Taxon Resolver interface illustrates automated resolution of historical or synonymous virus names to the currently accepted ICTV taxon, including lineage reconstruction and cross-references to external identifiers. The resolver is a reference implementation built on the JavaScript helper library that encapsulates Ontology Lookup Service (OLS) API calls. It accepts heterogeneous user inputs, including historical virus names, ICTV identifiers, Internationalized Resource Identifiers (IRIs), synonyms and NCBI Taxon identifiers, and applies a resolution strategy combining label matching, synonym detection, obsolete-term replacement chains and cross-taxonomy mappings. The output panel displays the currently accepted ICTV taxon together with its hierarchical lineage, rank, revision history and available cross-references to external resources. **Alt text:** Screenshot of the ICTV Taxon Resolver web interface showing a historical or synonymous virus taxon query and an output panel with the currently accepted ICTV taxon, lineage, rank, revision history and external cross-references.

Among the first operational deployments using the ICTV ontology API is the [EVORA Portal](https://portal.evora-project.eu/), a consolidated catalogue of research infrastructures’ resources and services in virology designed to support outbreak response and pandemic preparedness. The utilisation of the ICTV ontology API is integrated through one of the helper libraries directly into the catalogue generation workflow, enabling retrieval of up-to-date taxonomic terms along with associated lineage and synonym information. This ensures consistent use of the latest ICTV taxonomy, even when metadata provided by contributing research infrastructures correspond to earlier ICTV releases. As an added value for users of the research infrastructures, lineage information and historical synonyms retrieved through the ontology API enable enhanced search capabilities. This allows users to find resources using outdated taxa or alternative names, an option that was not available in the former search module.

Early adoption of the ICTV ontology API has also been observed in an external initiative concerned with pandemic preparedness and data integration: the [Pandemic PACT programme](https://www.pandemicpact.org/). This programme is now using ICTV taxonomy via the OLS API to align viral pathogen families and pathogens scientific names within its scope with the latest ICTV taxonomy (see Acknowledgements section). The retrieved ICTV metadata will be incorporated into [Pandemic PACT dashboard visualisations](https://www.pandemicpact.org/grants/visualise), where they will be integrated with metadata from other ontologies accessed through the OLS, such as the Systematized Nomenclature of Medicine (SNOMED), aligning up-to-date viral pathogen taxonomy with related information, including associated diseases.

Integration of ICTV taxonomy through an API enables timely and automated updates across resources. The [European Virus Archive (EVA)](https://www.european-virus-archive.com/), a network of distributed virus collections using a single web catalogue for virus and virus-related material ordering and distribution [18], plans to integrate the ICTV ontology into its workflows using OLS API queries to maintain local references to virus taxonomy aligned with the most recent ICTV releases. This will allow EVA datasets to remain synchronised with official taxonomy updates while preserving historical annotations associated with archived materials, which will enhance the interconnectivity of research and data management processes, improving resource discovery and data consistency.

These deployments demonstrate practical applicability across research infrastructures and public health platforms. The system enables automated resolution, cross-taxonomy linkage, reduced manual curation and reproducible versioning.

## 4 Discussion

Virus data resources include numerous databases with different scopes, data models, functionalities and degrees of FAIR maturity, as reflected in recent reviews of virus databases [19]. This heterogeneity creates practical challenges for interoperability and synchronisation with authoritative taxonomic references such as ICTV taxonomy. This work demonstrates how relational datasets can be easily transformed and exposed through scalable, public, ontology-based APIs. Standards-based ontology-enabled APIs drastically reduce the work needed to make datasets machine-actionable. Here, we developed an ontology for the widely used and regularly updated ICTV viral taxonomy and exposed it through an API, making it much easier to access, especially for automated update workflows. The ontology infrastructure acts as a semantic resolution layer that continuously bridges evolving authoritative ICTV releases with downstream computational ecosystems and legacy taxonomic references. Timely updates, in turn, reduce discrepancies that arise when downstream systems rely on outdated or partially synchronised taxonomic references and enable reproducible taxonomic resolution across computational workflows. Research infrastructures, sequence repositories, biodiversity catalogues, and knowledge bases can use automated alignment mechanisms to maintain consistency with official ICTV classifications, reducing the need for extensive manual curation.

Persistent ICTV identifiers and replacement relations provide a structured mechanism to track taxonomic evolution, historically prone to ambiguity and data fragmentation, in a computationally accessible way. The framework uses cross-references to build a pragmatic bridge between the authoritative, but annually updated, ICTV classification and widely used aggregation systems, such as NCBI taxonomy. This improves reproducibility and supports FAIR principles, and their progressive implementation [20, 21, 22]. While the framework substantially improves interoperability and historical resolution, synchronisation across downstream resources still depends on heterogeneous update cycles and integration strategies. In addition, some complex taxonomic transitions, such as splits, merges, or taxon abolitions, may still require complementary expert interpretation in specific downstream analytical contexts.

By exposing ICTV taxonomy through interoperable semantic technologies and a standardised API, the framework provides key interoperability components for connecting distributed data systems. Such capabilities are essential to FAIR-compliant federated environments, where independently maintained resources must remain discoverable, accessible and interoperable, including initiatives such as the European Open Science Cloud (EOSC) and the European Health Data Space (EHDS).

Consistent and machine-actionable taxonomy references support large-scale computational analyses, automated data integration workflows and reproducible comparative studies by reducing ambiguity in taxonomic interpretation. This contributes to improved retrieval and harmonisation of virological data across interoperable research infrastructures and data-sharing environments [23], supporting more reliable data integration, comparative analyses, evidence-based decision making, and more informed experimental design and data interpretation. The infrastructure also establishes a foundation for future enhancements, including expanded mappings with sequence repositories and the possible development of additional interfaces relying on the public OLS API.

Our approach aligns with the article “Four principles to establish a universal virus taxonomy” [24] that outlines key recommendations to achieve a coherent and comprehensive virus taxonomy. Our work helps make ICTV taxonomy more useful by ensuring that references to the official, evolution-based ICTV taxonomy, can be easily maintained and kept up-to-date, irrespective of the intended use or alternate classification scheme. For example, classifying viruses to support the surveillance and public health response to disease outbreaks ([Pathogens prioritization: a scientific framework for epidemic and pandemic research preparedness](https://www.who.int/publications/m/item/pathogens-prioritization-a-scientific-framework-for-epidemic-and-pandemic-research-preparedness)) results in a classification that is significantly different than that of the ICTV taxonomy. It is however still important to reference the current evolutionary-based classification so that our current knowledge on viruses can be quickly and easily adapted to newly identified or newly emerging ones. The ICTV taxonomy serves as the universal reference for virus classification, while the services we report here enable the critical links needed to connect that taxonomy to alternate schemes.

Adoption of the ICTV ontology API by resources relying on virus taxonomy (EVORA, and Pandemic PACT) demonstrates its practical applicability. Ontology-based APIs can facilitate alignment between biological taxonomy resources and clinical terminologies used in public health systems, enabling consistent pathogen identification across research infrastructures, data catalogues, and policy-oriented dashboards. By supporting automated retrieval of up-to-date taxonomic terms, lineage information and synonym relationships, the API-based approach supports consistent taxonomy usage across distributed data infrastructures and reduces the need for manual synchronisation of taxonomic references similarly to interoperability approaches explored in tools such as Treemendous for reconciling taxonomic backbones in tree species naming [25].

Taxonomic resources and biodiversity platforms such as NCBI Taxonomy and GBIF could use the OLS API to implement more automated synchronisation with ICTV releases. Databases such as ENA and GenBank, which depend on the NCBI taxonomy and its integration of ICTV classifications [26], would in turn benefit from improved data integration and accuracy. In cases where updates of viral taxonomy within NCBI do not immediately reflect the latest ICTV release, these resources could additionally leverage the ICTV ontology API, coupled with the provided SSSOM mappings, to maintain up-to-date viral taxonomy.

Our work builds on established FAIR and semantic standards [11, 27]. The inclusion of detailed metadata and the reuse of established semantic standards are critical for ensuring that the ICTV taxonomy serves as a reliable reference across different platforms and applications. This approach provides a lightweight and sustainable mechanism for publishing structured datasets as interoperable APIs, by leveraging an existing ontology service (OLS) and GitHub’s publicly available workflow automation. This approach is applicable to other domain-specific datasets. In the future, the ICTV plans to develop a dedicated API that will provide extended interoperability over a more comprehensive ICTV dataset, enhancing the OLS API and further ensuring FAIR compliance.

More broadly, this work illustrates how ontology technologies can expose structured datasets as semantically enriched graph representations through interoperable APIs. Traditional database schemas are optimised for storage and transactional operations but often require custom APIs to expose their content programmatically. In contrast, ontology-based representations allow relationships between entities to be explicitly encoded and navigated as a graph, enabling more flexible data integration and discovery. Such representations align with knowledge graph and Linked Open Data (LOD) approaches [28, 29]. More generally, this work demonstrates a reusable pattern for exposing structured, domain-specific datasets as interoperable ontologies and APIs by leveraging established ontology infrastructures, such as the OLS for biomedical ontologies.

## 5 Conclusion

This work shows that representing an evolving classification system, such as the ICTV taxonomy, as a unified ontology hosted through a public, scalable, standards-based API, increases its accessibility and addresses key challenges in access, versioning and interoperability. The resulting infrastructure supports backward-compatible resolution of viral taxonomy across historical ICTV releases, enabling historical traceability and alignment with current classifications through semantic modelling, persistent identifiers and interoperable mappings.

By building on open standards, automated workflows and existing ontology infrastructure services, this approach supports long-term sustainability and scalability while making ICTV taxonomy more readily integrable into computational workflows. It also facilitates synchronisation between authoritative taxonomic references and downstream biological, biomedical and public-health data infrastructures, supporting broader adoption of the latest official ICTV taxonomy as the reference standard and strengthening data consistency, reproducibility and preparedness for emerging viral threats, including through more timely dissemination of high-impact updates such as the pandemic-based designation of the species *Betacoronavirus pandemicum*.

Future developments may extend this framework toward broader interoperability with additional taxonomy providers, sequence repositories and interconnected semantic resources. More broadly, this work provides a reusable model for maintaining, exposing and reusing evolving taxonomic references through sustainable, machine-actionable semantic infrastructure.

## 6 Funding

This work was supported by the EVORA project (European Union’s HORIZON programme grant agreement N° 101131959). The contribution of the University of Alabama at Birmingham (UAB) team to the research reported in this publication was supported by the National Institute of Allergy and Infectious Diseases of the National Institutes of Health under Award Number [U24AI162625](https://reporter.nih.gov/search/pPQmLFOX30qEsVphAnsrIw/project-details/10269527). The content is solely the responsibility of the authors and does not necessarily represent the official views of the National Institutes of Health. The UAB team assisted with accessing all viral taxonomic data and associated metadata. The UAB team also served as consultants to assist in the understanding of the underlying ICTV data structures and data organization that were used to construct the ontology. In total, the UAB effort comprised approximately 10% of the overall effort.

## 7 Acknowledgements

This work is a product of the EVORA project (European Viral Outbreak Response Alliance) and the ICTV’s UAB team.

We gratefully acknowledge Emilia Antonio and Alice Norton from the Pandemic PACT programme for their feedback on the adoption of the ICTV OLS API within Pandemic PACT to align viral pathogen families and scientific names with the latest ICTV taxonomy identifiers.

## 8 Author contributions statement

Based on [Contributor Roles Taxonomy (CRediT)](https://credit.niso.org/)

- Conceptualization: PL
- Project administration: PL, BC
- Resources: CH, EL, PL, BC, JMCL, HP
- Software: JMCL, PL
- Methodology: PL, JMCL, HP, RD, CH
- Writing – original draft: PL, RD
- Writing – review & editing: PL, RD, HP, JMCL, BC, CH, EL
- Visualization: PL, RD, CH, DMD

How AI is used to help writing this paper:

- For rewording and style
- For tracking redundancy
- Ordinating arguments
- Checking forms considering the journal requirements and the final review (reference formats, link availability…)

## 9 Competing interests

The authors declare that they have no competing interests.

## 10 Data availability

All ontology artefacts, helper libraries, mapping resources, example notebooks, and continuous-integration workflows are publicly available under open licences. The ICTV MSL41 source data used in this work are available as the ICTVdatabase MSL41.v1 release (Hendrickson et al., 2026). The exact EVORA ICTV ontology release used for this manuscript is archived in Zenodo as EVORA-project/ictv-ontology: 2026-Apr-08-527a067 (McLaughlin, Lieutaud and Hendrickson, 2026). The exact ICTV-NCBI mapping release is archived in Zenodo as EVORA-project/virus-taxonomy-mappings: Virus Taxonomy Mappings Update, release-1 (McLaughlin and Lieutaud, 2026). Current development versions and latest releases are available from the public repositories listed below:

- ICTV database: https://github.com/ICTV-Virus-Knowledgebase/ICTVdatabase/
- ICTV ontology generation code, helper libraries, and generated ontologies (OWL/Turtle files): https://github.com/EVORA-project/ictv-ontology
- Latest ICTV ontology release: https://github.com/EVORA-project/ictv-ontology/releases/latest
- ICTV–NCBI SSSOM mappings: https://github.com/EVORA-project/virus-taxonomy-mappings
- Latest ICTV-NCBI mapping release: https://github.com/EVORA-project/virus-taxonomy-mappings/releases/latest
- Unified ICTV ontology in OLS: https://www.ebi.ac.uk/ols4/ontologies/ictv

All resources are reusable, openly licensed and versioned, enabling independent verification and downstream integration.

NCBI Taxonomy taxdump archives used to verify the integration status of ICTV releases are available from the NCBI taxonomy archive: https://ftp.ncbi.nih.gov/pub/taxonomy/taxdump_archive/

## 11 Code availability

- ICTV ontology generation code and helper libraries:
  - exact release archived at https://doi.org/10.5281/zenodo.20610329 [30]
  - current repository at https://github.com/EVORA-project/ictv-ontology
  - ICTV ontology API helper libraries at https://github.com/EVORA-project/ictv-ontology/tree/main/helpers
- ICTV-NCBI mapping generation:
  - exact release archived at https://doi.org/10.5281/zenodo.20610593 [31]
  - current repository at https://github.com/EVORA-project/virus-taxonomy-mappings
- ICTV Taxon Resolver source code: https://github.com/EVORA-project/ictv-resolver
- ICTV Taxon Resolver live public instance: https://evora-project.github.io/ictv-resolver
- EMBL-EBI Ontology Lookup Service (OLS): https://github.com/EBISPOT/ols4/blob/dev/README.md

## 12 Acronyms

API(s): Application Programming Interface(s)
CI/CD: Continuous Integration and Deployment
COL: Catalogue of Life
CURIE: Compact Uniform Resource Identifier
DCTERMS: Dublin Core Terms namespace
ENA: European Nucleotide Archive
ETL process: Extract, Transform, and Load
EVA: European Virus Archive
EVORA: European Viral Outbreak Response Alliance.
FAIR: Findable, Accessible, Interoperable and Reusable
FOAF: Friend of a Friend ontology
GBIF: Global Biodiversity Information Facility
IAO: Information Artifact Ontology
ICTV: International Committee on Taxonomy of Viruses
IRI: Internationalized Resource Identifier
LOD: Linked Open Data
MCP: Model Context Protocol
MSL: Master Species List
NCBI: National Center for Biotechnology Information
OLS: Ontology Lookup Service
OWL: Web Ontology Language
PROV-O: Provenance Ontology
RDFS: Resource Description Framework Schema
REST: Representational State Transfer
SKOS: Simple Knowledge Organization System
SNOMED: Systematized Nomenclature of Medicine
SSSOM: Simple Standard for Sharing Ontology Mapping
TAXRANK: Taxonomic Rank Vocabulary
VMR: Virus Metadata Resource

## Notes

https://github.com/EVORA-project/ictv-ontology

https://github.com/EVORA-project/ictv-ontology/releases/latest

https://github.com/EVORA-project/virus-taxonomy-mappings

https://github.com/EVORA-project/virus-taxonomy-mappings/releases/latest

https://github.com/ICTV-Virus-Knowledgebase/ICTVdatabase/

### Competing Interest Statement

The authors have declared no competing interest.

## References

1. Zerbini FM, Siddell SG, Mushegian AR, Walker PJ, Lefkowitz EJ, et al. Differentiating between viruses and virus species by writing their names correctly. Arch Virol 2022;167:1231–1234. doi:10.1007/s00705-021-05323-4

2. Simmonds P, Adriaenssens EM, Lefkowitz EJ et al. Changes to virus taxonomy, the international code of virus classification and nomenclature, and the ICTV statutes ratified by the International Committee on Taxonomy of Viruses (2025). Arch Virol 2026;171:23. doi:10.1007/s00705-025-06485-1

3. Black EJ, et al. Virus taxonomy: the database of the International Committee on Taxonomy of Viruses. Nucleic Acids Res 2026;54(D1). doi:10.1093/nar/gkaf1159

4. Boyle B, Hopkins N, Lu Z, et al. The taxonomic name resolution service: an online tool for automated standardization of plant names. BMC Bioinformatics 2013;14:16. doi:10.1186/1471-2105-14-16

5. Malone J, Stevens R, et al. Ten simple rules for selecting a bio-ontology. PLoS Comput Biol 2016;12(2):e1004743. doi:10.1371/journal.pcbi.1004743

6. McMurry JA, Juty N, Blomberg N, et al. Identifiers for the 21st century: how to design, provision, and reuse persistent identifiers. PLoS Biol 2017;15(6):e2001414. doi:10.1371/journal.pbio.2001414

7. Hendrickson RC, Mims L, Lefkowitz EJ. ICTV-Virus-Knowledgebase/ICTVdatabase: Taxonomy Release: MSL41.v1 20260320 [Dataset]. Zenodo 2026. doi:10.5281/zenodo.19339543

8. Kibbe WA, Arze C, Felix V, et al. Disease Ontology 2015 update: an expanded and updated database of human diseases for linking biomedical knowledge through disease data. Nucleic Acids Res 2015;43 (Database issue). doi:10.1093/nar/gku1011

9. Lebo T, Sahoo S, McGuinness D, et al. PROV-O: the PROV ontology. W3C Recommendation 2013. Available at: https://www.w3.org/TR/prov-o/ (Accessed: 12 May 2026)

10. Miles A, Bechhofer S. SKOS simple knowledge organization system reference. W3C Recommendation 2009. Available at: https://www.w3.org/TR/skos-reference/ (Accessed: 12 May 2026)

11. Xu F, Juty N, Goble C, Jupp S, Parkinson H, Courtot M. Features of a FAIR vocabulary. J Biomed Semantics 2023;14(1):6. doi:10.1186/s13326-023-00286-8

12. Smith B, Ashburner M, Rosse C, et al. The OBO Foundry: coordinated evolution of ontologies to support biomedical data integration. Nat Biotechnol 2007;25(11):1251–1255. doi:10.1038/nbt1346

13. Fielding RT, Taylor RN. Principled design of the modern Web architecture. ACM Trans Internet Technol 2002;2(2):115–150. doi:10.1145/337180.337228

14. McLaughlin J, Lagrimas J, Iqbal H, et al. OLS4: a new Ontology Lookup Service for a growing interdisciplinary knowledge ecosystem. Bioinformatics 2025;41(5):btaf279. doi:10.1093/bioinformatics/btaf279

15. David R, Rybina A, Burel J, et al. “Be sustainable”: EOSC-Life recommendations for implementation of FAIR principles in life science data handling. EMBO J 2023a;42:e115008. doi:10.15252/embj.2023115008

16. David R, Baumann K, LeFranc Y, et al. Converging on a semantic interoperability framework for the European Data Space for Science, Research and Innovation (EOSC). In: Proceedings of the 2nd Workshop on Ontologies for FAIR and FAIR Ontologies (Onto4FAIR). 2023b. doi:10.5281/zenodo.8102786

17. Matentzoglu N, Balhoff J, Bello S, et al. A simple standard for sharing ontological mappings (SSSOM). Database 2022;2022:baac035. doi:10.1093/database/baac035

18. Romette JL, Prat CM, Gould EA, et al. The European Virus Archive goes global: a growing resource for research. Antiviral Res 2018;158:127–134. doi:10.1016/j.antiviral.2018.07.017

19. Ritsch M, Cassman NA, Saghaei S, Marz M. Navigating the Landscape: A Comprehensive Review of Current Virus Databases. Viruses 2023;15(9):1834. doi:10.3390/v15091834

20. Wilkinson MD, Dumontier M, Aalbersberg IJ, et al. The FAIR guiding principles for scientific data management and stewardship. Sci Data 2016;3:160018. doi:10.1038/sdata.2016.18

21. Sansone SA, McQuilton P, Rocca-Serra P, et al. FAIRsharing as a community approach to standards, repositories and policies. Nat Biotechnol 2019;37(4):358–367. doi:10.1038/s41587-019-0080-8

22. David R, Mabile L, Specht A, et al. FAIRness literacy: the Achilles’ heel of applying FAIR principles. Data Sci J 2020;19(1):32. doi:10.5334/dsj-2020-032

23. Research Data Alliance COVID-19 Working Group. Recommendations and guidelines on data sharing. 2020. doi:10.15497/rda00052

24. Simmonds P, Adriaenssens EM, Zerbini FM, et al. Four principles to establish a universal virus taxonomy. PLoS Biol 2023;21(2):e3001922. doi:10.1371/journal.pbio.3001922

25. Specker F, Paz A, Crowther TW, Maynard DS. Treemendous: an R package for integrating taxonomic information across backbones. PeerJ 2024;12:e16896. doi:10.7717/peerj.16896.

26. Schoch CL, Ciufo S, Domrachev M, et al. NCBI taxonomy: a comprehensive update on curation, resources and tools. Database 2020;2020:baaa062. doi:10.1093/database/baaa062

27. Berners-Lee T. Linked Data. 2006. Available at: https://www.w3.org/DesignIssues/LinkedData.html (Accessed: 09 June 2026)

28. Hogan A, Blomqvist E, Cochez M, et al. Knowledge graphs. ACM Comput Surv 2021;54(4):1–37. doi:10.1145/3447772

29. Kamdar MR, Musen MA. An empirical meta-analysis of the life sciences linked open data on the web. Sci Data 2021;8:24. doi:10.1038/s41597-021-00797-y

30. McLaughlin J, Lieutaud P, Hendrickson C. EVORA-project/ictv-ontology: 2026-Apr-08-527a067 [Software]. Zenodo 2026. doi:10.5281/zenodo.20610329

31. McLaughlin J, Lieutaud P. EVORA-project/virus-taxonomy-mappings: Virus Taxonomy Mappings Update [Software]. Zenodo 2026. doi:10.5281/zenodo.20610593

## Web resources

Catalogue of Life (COL). Available at: https://www.catalogueoflife.org/ (accessed: 09 June 2026).

Catalogue of Life ICTV Master Species List dataset. Available at: https://www.catalogueoflife.org/data/dataset/1014 (accessed: 09 June 2026).

Contributor Roles Taxonomy (CRediT). Available at: https://credit.niso.org/ (accessed: 09 June 2026).

EMBL-EBI Ontology Lookup Service (OLS4). Available at: https://www.ebi.ac.uk/ols4/index (accessed 09 June 2026).

EMBL-EBI OLS4 API documentation. Available at: https://www.ebi.ac.uk/ols4/api-docs (accessed 09 June 2026).

EMBL-EBI OLS4 server documentation. Available at: https://github.com/EBISPOT/ols4/blob/dev/README.md (accessed 09 June 2026).

European Nucleotide Archive (ENA). Available at: https://www.ebi.ac.uk/ena/browser/home (accessed: 09 June 2026).

European Viral Outbreak Response Alliance (EVORA). Available at: https://evora-project.eu/ (accessed: 09 June 2026).

European Viral Outbreak Response Alliance (EVORA) project description at CORDIS. Available at: 10.3030/101131959 (accessed: 09 June 2026).

EVORA ICTV ontology helper and API documentation. Available at: https://evora-project.github.io/ictv-ontology/helpers/ (accessed: 09 June 2026).

EVORA ICTV ontology GitHub repository. Available at: https://github.com/EVORA-project/ictv-ontology (accessed 09 June 2026).

EVORA ICTV helper libraries GitHub directory. Available at: https://github.com/EVORA-project/ictv-ontology/tree/main/helpers (accessed 09 June 2026).

EVORA ICTV Taxon Resolver live interface. Available at: https://evora-project.github.io/ictv-resolver (accessed 09 June 2026).

EVORA ICTV Taxon Resolver GitHub repository. Available at: https://github.com/EVORA-project/ictv-resolver (accessed 09 June 2026).

EVORA Portal. Available at: https://portal.evora-project.eu/ (accessed: 09 June 2026).

EVORA virus-taxonomy-mappings GitHub repository. Available at: https://github.com/EVORA-project/virus-taxonomy-mappings (accessed 09 June 2026).

European Virus Archive (EVA). Available at: https://www.european-virus-archive.com/ (accessed: 09 June 2026).

GenBank. Available at https://www.ncbi.nlm.nih.gov/genbank/ (accessed: 09 June 2026).

Global Biodiversity Information Facility (GBIF). Available at: https://www.gbif.org/ (accessed: 09 June 2026).

ICTV Master Species List (MSL). Available at: https://ictv.global/msl (accessed 09 June 2026).

ICTV Virus Metadata Resource (VMR). Available at: https://ictv.global/vmr (accessed 09 June 2026).

ICTV Virus Knowledgebase GitHub repository. Available at: https://github.com/ICTV-Virus-Knowledgebase (accessed: 09 June 2026).

ICTV database GitHub repository. Available at: https://github.com/ICTV-Virus-Knowledgebase/ICTVdatabase/ (accessed 09 June 2026).

International Committee on Taxonomy of Viruses (ICTV). Available at: https://ictv.global/ (accessed: 09 June 2026).

International Union of Microbiological Societies (IUMS). Available at: https://www.iums.org/ (accessed 09 June 2026).

National Center for Biotechnology Information (NCBI). Available at: https://www.ncbi.nlm.nih.gov/ (accessed: 09 June 2026).

NCBI taxonomy browser. Available at: https://www.ncbi.nlm.nih.gov/datasets/taxonomy/browser/ (accessed: 09 June 2026).

NCBI taxonomy archive (taxdump archive). Available at: https://ftp.ncbi.nih.gov/pub/taxonomy/taxdump_archive/ (accessed: 09 June 2026).

Pandemic PACT. Available at: https://www.pandemicpact.org/ (accessed: 09 June 2026).

Pandemic PACT dashboard visualisations. Available at: https://www.pandemicpact.org/grants/visualise (accessed: 09 June 2026).

Simple Standard for Sharing Ontological Mappings (SSSOM) standard documentation. Available at: https://mapping-commons.github.io/sssom/1.0/ (accessed: 09 June 2026).

Unified ICTV ontology in OLS. Available at: https://www.ebi.ac.uk/ols4/ontologies/ictv (accessed 09 June 2026).

ViralZone. Available at: https://viralzone.expasy.org/ (accessed: 09 June 2026).

Virus Host DB. Available at: https://www.genome.jp/virushostdb/ (accessed: 09 June 2026).

WHO Pathogens prioritization: a scientific framework for epidemic and pandemic research preparedness. Available at: https://www.who.int/publications/m/item/pathogens-prioritization-a-scientific-framework-for-epidemic-and-pandemic-research-preparedness (accessed: 09 June 2026).

Wikidata. Available at: https://www.wikidata.org/ (accessed: 09 June 2026).

