## Supplementary Material for "Programmatic access to ICTV virus taxonomy through a public ontology API"

**Abstract**

This file contains supplementary material supporting the main manuscript, including API usage examples, helper library architecture, the taxonomy-provider landscape, and the ICTV-NCBI mapping specification.

### 1 Supplementary Material

This supplementary file accompanies the manuscript and provides implementation-oriented documentation, developer guidance, and additional context for comparative evaluation and mapping specifications.

#### 1.1 Supplementary Material S1 – API usage examples

In addition to the generic OLS4 API documentation, a dedicated, ICTV-specific documentation page is provided on EVORA ICTV-ontology GitHub repository. This resource documents both direct use of the ICTV ontology via the generic OLS API, and use of the ICTV-specific helper libraries (see Supplementary Material S2), and enables implementation of common taxonomic resolution workflows. It complements the general OLS documentation by specifically addressing ICTV-related use cases:

- OLS4 Server documentation: `https://github.com/EBISPOT/ols4/blob/dev/README.md`
- OLS4 API documentation: `https://www.ebi.ac.uk/ols4/api-docs`
- EVORA ICTV helper and API documentation page: `https://evora-project.github.io/ictv-ontology/helpers/`

Typical EVORA/ICTV use cases include retrieving taxonomy entities, resolving obsolete taxa, reconstructing lineage information, and retrieving synonyms and historical names.

OLS4 provides multiple REST controllers for querying classes, properties, entities, individuals, search, and autosuggestion, with two stable entry points: v1 controllers (legacy endpoints, including useful features such as /api/suggest ) and v2 controllers (more recent endpoints with richer metadata on linked entities, particularly suited for taxon resolution).

Together, these endpoints support typical ICTV use cases. For example, the paginated v1 /terms endpoint provides access to the full ontology, including historical and obsolete terms, whereas the v2 /classes endpoint enables more targeted retrieval of taxa through search strategies focusing on specific fields. Historical names and obsolete taxa can be resolved using metadata integrated into the ontology, notably oboInOwl:hasExactSynonym and IAO:0100001 (term_replaced_by) from Information Artifact Ontology (IAO).

As an example, resolving the currently accepted taxon name corresponding to the species formerly known as "*Severe acute respiratory syndrome coronavirus*" requires retrieving historical occurrences among obsolete entities, and then following the IAO:0100001 relation to the current replacement taxon.

Stable ICTV identifiers, such as ICTV20040588, serve as direct entry points. The EVORA repository further provides example notebooks and utilities illustrating typical workflows such as querying taxa by label or ICTV identifier, following obsolescence relationships, and reconstructing the history of a taxon across releases.

### 1.2 Supplementary Material S2 – Helper library architecture

Helper libraries, developed in JavaScript, Python and PHP, simplify integration into computational pipelines. These libraries encapsulate the main ICTV resolution use cases logic on top of the OLS API and hide the complexity of direct ontology queries from downstream developers.

The JavaScript helper is provided as a browser-compatible JavaScript module for web applications. It exposes methods for resolving terms to the latest accepted ICTV taxon, retrieving taxon revision histories across MSL releases, resolving virus-name individuals through their parent taxa, handling direct taxon names and ICTV or NCBI Taxon identifiers, and returning lineage information.

The PHP helper provides equivalent functionality for PHP-based services, allowing resolution of identifiers and names, traversal of replacement chains, export of releases and retrieval of isolate-related information.

The Python helper provides the same high-level operations for scripts, Extract, Transform, and Load (ETL) workflows and Jupyter-based analyses. This makes it suitable for pipeline integration and reproducible computational notebooks.

Across all three implementations, the same general resolution strategy is applied:

1. recognise ICTV IRIs and ICTV IDs;

2. optionally map NCBI Taxon identifiers to ICTV taxa;

3. treat individuals, such as isolates, as entry points and resolve them to the corresponding species;

4. fall back to labels and synonyms where needed;

5. when an entity is obsolete, follow the replacement chain until final non-obsolete taxa are reached.

The libraries return a normalised ICTV taxon object containing identifiers, IRI, label, synonyms, rank, lineage, obsolescence status, revision links and, where available, NCBI mappings. This abstraction allows developers to focus on domain logic rather than the details of the OLS API.

Description examples and further details available from:
`https://github.com/EVORA-project/ictv-ontology/tree/main/helpers`

A public browser-based ICTV Taxon Resolver was built as a reference implementation of this logic using the JavaScript helper library. It accepts historical or current virus names, ICTV identifiers,

IRIs and NCBI Taxon identifiers, and returns the latest accepted ICTV taxon together with its lineage and mapping information.

To demonstrate how these helpers can be used in practice, we built a public, browser-based ICTV Taxon Resolver, using the JavaScript library. It is available as open source here:

`https://github.com/EVORA-project/ictv-resolver`

with a live, interactive instance at:

`https://evora-project.github.io/ictv-resolver`

This tool serves both as a practical service for virologists and as a didactic example of how to integrate the OLS-backed ICTV ontology into web applications such as those used by EVORA.

### 1.3 Supplementary Material S3 – Landscape of viral taxonomy providers and integration patterns (June 2026)

This supplementary section provides a qualitative overview of major resources involved in the dissemination, integration, and reuse of viral taxonomy. This analysis aims to characterise how ICTV taxonomy is propagated across the data resource ecosystem and to highlight practical implications for programmatic use.

The comparison focuses on key dimensions relevant to computational workflows, including: (i) recency relative to ICTV releases, (ii) availability of historical names and synonym information, (iii) persistence of identifiers across releases, (iv) availability of programmatic access (e.g. APIs or bulk downloads), (v) support for lineage reconstruction, and (vi) integration with external resources.

#### 1.3.1 Authoritative source

**International Committee on Taxonomy of Viruses (ICTV)** ICTV is the official authority responsible for virus classification and nomenclature. It publishes the Master Species List (MSL) and associated data files, which constitute the reference taxonomy used across virology, as well as the Virus Metadata Resource (VMR) which links the species in that reference taxonomy to exemplar virus isolate genomes in GenBank. At the time of submission of this manuscript, the most recent ICTV release is MSL41.v1, with 22 670 taxonomic terms, released in March 2026.

ICTV resources are comprehensive and regularly updated; they provide access to historical information through structured files and web interfaces. However, access to this information is primarily designed for human use. While raw data files are publicly available, no public API has historically been provided to support programmatic access to taxonomic history, automated resolution of obsolete terms, or identifier-based queries across releases. As a result, integration into computational workflows requires additional processing and interpretation.

#### 1.3.2 Integration platforms embedding ICTV taxonomy

**NCBI Taxonomy** NCBI Taxonomy provides a widely used taxonomic backbone integrated with sequence databases such as GenBank and ENA. It incorporates ICTV taxonomy and offers persistent identifiers, extensive cross-links to biological data, and mature programmatic access via the E-utilities APIs and bulk downloads. In addition, NCBI provides monthly historical taxdump archives as dated files named new_taxdump_YYYY-MM-01.zip, which allow changes in NCBI taxonomic content to be retrospectively examined. The observations described below were checked against the April, May and June 2026 archives (new_taxdump_2026-04-01.zip, new_taxdump_2026-05-01.zip and new_taxdump_2026-06-01.zip in the taxdump archives), the first three monthly archives after the March 2026 MSL41 release, and against the live NCBI Taxonomy service on 10 June 2026.

However, integration of ICTV releases has historically lagged behind the most recent official version and sometimes occurs progressively. As of April 2026 archive, NCBI appeared to still reflect the previous (MSL40) release for the examples checked: the recently (MSL41) renamed species "*Mamastrovirus suisorientalis*" (ICTV19950506) was absent, although the corresponding NCBI taxon later shown as ncbitaxon:3430637 was represented under its previous name, and the newly created species "*Mastadenovirus himalaiense*" (ICTV202522146) was absent from NCBI Taxonomy. In May 2026, renamed taxa such as "*Mamastrovirus suisorientalis*" (ICTV19950506, ncbitaxon:3430637) had begun to appear, while in the June 2026 archive and in the live NCBI Taxonomy service checked on 10 June 2026, the newly created taxa checked here were still absent, suggesting a progressive integration of the latest release (MSL41).

In the June 2026 archive, and in the live NCBI Taxonomy service checked on 10 June 2026, the abolished ICTV taxa could also still be found in NCBI Taxonomy with taxonomic ranks and labelled as ICTV-accepted names. Examples include the abolished order "*Petitvirales*" (ICTV201907169, ncbitaxon:2732414), still displayed as an order, and the abolished species "*Nepovirus americaense*" (ICTV19911709, ncbitaxon:3047734), still displayed as a species.

In addition, NCBI maintains historical and intermediate taxonomic states to preserve consistency with sequence records. While this approach is essential for data stability, it may introduce discrepancies with the latest ICTV classification and complicate resolution of current accepted names or taxonomy. Furthermore, extensions below ICTV-defined ranks may introduce additional heterogeneity relative to the official taxonomy. As an illustrative example, the former ICTV species taxon name "*Severe acute respiratory syndrome-related coronavirus*" still has an entry conserving this exact name at NCBI known as ncbitaxon:694009, representing an NCBI-specific entry rather than a current ICTV-accepted taxon.

**Catalogue of Life and GBIF** The Catalogue of Life (COL) and the Global Biodiversity Information Facility (GBIF) provide broad biodiversity integration and global dissemination infrastructures. Both incorporate viral taxonomy derived from ICTV releases, either directly (COL) or through intermediary providers (GBIF via COL). As of June 2026 COL still integrates ICTV MSL38 and is three full releases behind ICTV (source: `https://www.catalogueoflife.org/data/dataset/1014` accessed 3 June 2026).

These platforms offer programmatic access and facilitate interoperability across biodiversity datasets. However, their update cycles are independent of ICTV, which may result in delays relative to the latest official taxonomy. In addition, their data models generally prioritise current classifications and may provide limited support for explicit representation of historical terms or systematic resolution of obsolete names. When intermediary providers are involved, additional latency and transformation layers may further affect consistency with the source taxonomy.

#### 1.3.3 Downstream consumers of viral taxonomy

Resources such as ViralZone, EVA (European Virus Archive), GenBank, ENA, and Virus-Host DB incorporate viral taxonomic information primarily for data organisation, annotation, and discovery. These systems either integrate ICTV taxonomy directly from source files or rely on other providers (e.g. NCBI Taxonomy or other intermediaries) for classification. As a result, their taxonomic representations depend both on their own update cycles and on those of their upstream providers, along with the associated limitations. While they provide essential domain-specific functionality, they generally do not aim to expose the full structure, history, or evolution of viral taxonomy in a form suitable for programmatic reconciliation across releases.

A distinct case is Wikidata, which integrates and links heterogeneous data from multiple sources within a knowledge graph framework. While this enables rich cross-resource connections, taxonomic entities originating from different sources or time points may coexist without explicit, machine readable links reflecting their historical relationships. For example, Wikidata may contain both the former ICTV taxon species "Zika virus" (ICTV19990862, wikidata:Q202864) and the current corresponding ICTV taxon species "Orthoflavivirus zikaense" (ICTV19990862, wikidata:Q118455511) as separate entities representing the same species, without indicating deprecation or providing a systematic mechanism to resolve their taxonomic continuity.

#### 1.3.4 Synthesis and implications

This analysis highlights that, although ICTV taxonomy is widely disseminated and reused across multiple platforms, significant heterogeneity remains in how it is represented, updated, harmonised and accessed in data resources [1]. In particular, differences are observed in update latency relative to ICTV releases, persistence and interpretation of identifiers, and support for historical term resolution.

No single resource currently provides a unified solution combining:
(i) up-to-date alignment with ICTV releases,
(ii) explicit and machine-actionable representation of taxonomic history, and
(iii) standardised programmatic access enabling automated resolution of obsolete or historical terms.

The ontology-based infrastructure presented in this work addresses these limitations by providing a unified, semantically explicit, and historically traceable representation of ICTV taxonomy, together with a programmatic access layer through the OLS API. This approach enables consistent resolution of taxonomic entities across releases and facilitates integration of ICTV taxonomy into computational workflows.

### 1.4 Supplementary Material S4 – ICTV–NCBI mapping specification

Interoperability between ICTV and NCBI taxonomy is supported through an ICTV–NCBI mapping compiled and published, in SSSOM format (`https://mapping-commons.github.io/sssom/1.0/`) in the EVORA virus-taxonomy-mappings repository:

`https://github.com/EVORA-project/virus-taxonomy-mappings`

The mapping file (ictv_ncbitaxon_exact.sssom.tsv) contains exact lexical matches between ICTV taxa, represented as ICTV CURIEs, and corresponding NCBI Taxon identifiers. The file header declares the CURIE map, license, mapping provider, mapping set identifier, and mapping set title. At row level, the mapping table includes the subject and object identifiers and labels, the predicate, the mapping justification, the mapping tool/provider, and the mapping date.

Mappings are expressed using skos:exactMatch [2], and the justification is indicated as semapv:LexicalMatching. The repository URL is identified as the mapping provider tool that generated the mapping, and each mapping row includes a mapping date. These elements provide basic mapping metadata and document the origin and method of the generated correspondences.

A mapping layer in the helper libraries (see Supplementary Material S2) loads this SSSOM file and supports both ICTV → NCBI and NCBI → ICTV lookups. This enables downstream resources to align locally stored NCBI-based annotations with official ICTV taxonomy and *vice versa*.

Because the mappings are expressed in the SSSOM standard, they can also be reused by generic ontology-mapping tooling and broader cross-mapping services, strengthening interoperability across the life-science ontology ecosystem.

### 2 References


1. Seah BKB. Paying it forward: crowdsourcing the harmonisation and linking of taxon names and biodiversity identifiers. Biodivers Data J 2023;11:e114076. doi:10.3897/BDJ.11.e114076

2. Miles A, Bechhofer S. SKOS simple knowledge organization system reference. W3C Recommendation 2009. Available at: https://www.w3.org/TR/skos-reference/ (Accessed: 12 May 2026)
